# Effects of temperature and *p*CO_2_ on the respiration, biomineralization and photophysiology of the giant clam *Tridacna maxima*

**DOI:** 10.1101/672907

**Authors:** Chloé Brahmi, Leila Chapron, Gilles Le Moullac, Claude Soyez, Benoît Beliaeff, Claire E. Lazareth, Nabila Gaertner-Mazouni, Jeremie Vidal-Dupiol

## Abstract

Such as many other reef organisms, giant clams are today confronted to global change effects and can suffer mass bleaching or mortality events mainly related to abnormally high seawater temperatures. Despite its strong ecological and socio-economical importance, its responses to the two most alarming threats linked to global change (i.e., ocean warming and acidification) still need to be explored. We investigated physiological responses of 4-years-old *Tridacna maxima* specimens to realistic levels of temperature and partial pressure of carbon dioxide (*p*CO_2_) (+1.5°C and +800 *μ*atm of CO_2_) predicted for 2100 in French Polynesian lagoons during the warmer season. During a 65-days crossed-factor experiment, individuals were exposed to two temperatures (29.2°C; 30.7°C) and two *p*CO_2_ (430 *µ*atm; 1212 *µ*atm) conditions. Impact of each parameter and their potential synergetic effect were evaluated on respiration, biomineralization and photophysiology. Kinetics of thermal and acidification stress were evaluated by performing measurements at different times of exposure (29, 41, 53, 65 days). At 30.7°C, the holobiont O_2_ production, symbiont photosynthetic yield, and density were negatively impacted. High *p*CO_2_ had a significant negative effect on shell growth rate, symbiont photosynthetic yield and density. Shell microstructural modifications were observed from 41 days in all temperature and *p*CO_2_ conditions. No significant synergetic effect was found. Today thermal conditions (29.2°C) appeared to be sufficiently stressful to induce a host acclimatization process. All these observations indicate that temperature and *p*CO_2_ are both forcing variables affecting *T. maxima* physiology and jeopardize its survival under environmental conditions predicted for the end of this century.

## 1. Introduction

Since the industrial revolution, human activities released gigatons of CO_2_ in the atmosphere, which participate to global change (Broeker et al., 1979; Caldeira and Wickett, 2003; Sabine et al., 2004; Zeebe et al., 2008; IPCC, 2014). Due to the “greenhouse gas” property of CO_2_, terrestrial and sea surface temperature rise constantly (i.e., global warming). According to the last Intergovernmental Panel on Climate Change report (IPCC, 2018), human-induced warming had already reached about +1°C above pre-industrial levels and the panel predicts +1.5°C around 2040 if the current warming rate continues. Enrichment of dissolved CO_2_ in the ocean modifies the carbonate chemistry by depressing carbonate ion concentration ([CO_3_^2-^]) and releasing protons which respectively decrease calcium carbonate saturation state (Ω) and seawater pH (Kleypas et al., 1999; Caldeira and Wickett, 2003) (i.e., ocean acidification). Indeed, pH had already decreased by 0.1 pH unit since the pre-industrial revolution and the decrease may reach −0.3 to −0.4 pH unit by 2100 (according to the median scenario RCP 4.5, IPCC, 2014).

Since CO_3_^2-^ is a carbonate ion form involved in biologically controlled calcification process (i.e., biomineralization) (Orr et al., 2005), marine organisms which form calcium carbonate structures (e.g., exoskeleton, shell, test, spicule), such as scleractinian corals, mollusks, and echinoderms are particularly vulnerable to ocean acidification (Hoegh-Guldberg et al., 2007; Ries et al., 2009; Kroeker et al., 2013; Gazeau et al., 2013). Among marine calcifying mollusks, negative impacts of temperature and *p*CO_2_ have been demonstrated on the survival, growth, biomineralization processes, and others key physiological functions on different stages of their life cycle (Bougrier et al., 1995; Kurihara, 2008; Gazeau et al., 2010, 2011; Schwartzmann et al., 2011; Rodolfo-Metalpa et al., 2011; Talmage and Gobler, 2011; Watson et al., 2012; Liu et al., 2012; Kuhihara et al., 2013; Fitzer et al., 2014; Le Moullac et al., 2016a,b; Wessel et al., 2018). However, effects of each stressor and their potential synergetic effect on giant clam physiology are still poorly known.

Giant clams live in obligatory symbiosis with photosynthetic dinoflagellates of the family Symbiodiniaceae (Holt et al., 2014; LaJeunesse et al., 2018). Different giant clam species, i.e., *Tridacna maxima, Tridacna crocea, Tridacna noae* and *Tridacna squamosa*, have been found to associate with different Symbiodiniaceae genera, i.e., *Symbiodinium* (formerly clade A, LaJeunesse et al., 2018), *Cladocopium* (formerly clade C, LaJeunesse et al., 2018) and *Durusdinium* (formerly clade D, LaJeunesse et al., 2018) (Pinzón et al., 2011; DeBoer et al., 2012; Ikeda et al., 2017; Lim et al., 2019). Hereafter, the symbiotic Symbiodiniaceae are named “symbionts”. In giant clams, symbionts are located extracellularly inside a tubular system called “Z-tubules” found in the outer epithelium of the mantle and connected to the stomach (Norton et al., 1992; Holt et al., 2014). These mixotrophic organisms acquire nutrients from heterotrophic (via seawater filtration) and symbionts photo-autotrophic pathways (Klumpp et al., 1992; Hawkins and Klumpp, 1995). Nutrients provided by symbionts (such as glucose, Ishikura et al., 1999) may account for a major part of the clam energy needs depending on the species and life stage (Trench et al., 1981; Klumpp et al., 1992; Klumpp and Lucas, 1994; Klumpp and Griffits, 1994; Hawkins and Klumpp, 1995; Elfwing et al., 2002; Yau and Fan, 2012; Holt et al., 2014; Soo and Todd, 2014). Like symbiotic coral species, bleaching events affecting giant clam have been recorded several times in the Indo-Pacific Ocean (Adessi, 2001; Buck et al., 2002; Leggat et al., 2003; Andréfouët et al., 2013; Junchompoo et al., 2013) mainly due to seawater temperature increase by a few degrees above the seasonal maximum (Adessi, 2001; Andréfouët et al., 2013). Bleaching process results from the symbiosis breakdown via the loose of symbiotic Symbiodiniaceae (Buck et al., 2002; Leggat et al., 2003).

In French Polynesia, *Tridacna maxima* species (Röding, 1798) is one of the most emblematic and patrimonial organisms. They represent an important food resource for inhabitants of remote atolls and giant clam fishery and sustainable aquaculture activities generate substantial incomes for local fishermen and farmers (Van Wysberge et al., 2016; Andréfouët et al., 2017). However, wild and cultivated giant clam stocks are largely threatened by environmental disturbances such as abnormally high sea surface temperature inducing mass mortality events as reported in Tuamotu atolls by Adessi (2001) and Andréfouët et al. (2013). More recently, two mass bleaching events occurring in Reao and Tatakoto (Tuamotu islands) were linked to a prolonged high lagoonal temperature (≥ 30°C over several weeks) of these semi-closed atolls (Andréfouet et al., 2017; Fig. S1). All these observations strongly indicate that temperature is an important stressor for giant clams. Besides the ecological consequences, giant clam mass bleaching and mortality cause a significant loss of incomes for the islanders.

Despite their ecological and socio-economic importance, effects of thermal stress and acidification on giant clams still need to be investigated. Thermal stress has shown to decrease fertilization success (Amstrong, 2017) in *Tridacna maxima* and oxygen production and respiration rates in *T. gigas* and *T. deresa* species (Blidberg et al., 2000). In contrast, an increase of respiration with a high photosynthetic rate in the holobiont was reported for *T. squamosa* (Elfwing et al., 2001). Thermal stress also reduces the abundance of symbionts in *T. gigas* (Leggat et al., 2003) even after only 12h of exposure in *T. crocea* (Zhou et al., 2018) and a decrease of photosynthate export from symbionts to the host (Leggat et al., 2003). In *T. maxima*, heat stress induced changes of fatty acid composition and lipid pathways and ROS (Reactive Oxygen Species) scavenger overexpression (Dubousquet et al., 2016). Combined to high light intensities, heat stress caused a decrease of the cell size of remaining symbionts and of their chlorophyll content (Buck et al., 2002). Recent studies demonstrated that low pH seawater with high nutrient concentration could have variable effects on growth on four species of *Tridacna* genus (Toonen et al., 2012). Moreover, Watson (2015) showed that high light irradiance condition limits the impact of *p*CO_2_ on shell growth. Regarding the potential synergetic effect of temperature and *p*CO_2_ parameters, the only study ever carried out showed that ocean warming and acidification may reduce survival in *T. squamosa* juveniles (Waston et al., 2012). To our knowledge, no study had already investigated impact of both parameters, and their potential synergetic effect, on several key physiological parameters of both *Tridacna maxima* host and its symbionts. To fill this gap, our experimental approach was designed to better understand the physiological mechanisms underlying the response of giant clams to global climate changes.

Based on a 65 days crossed-factor experiment, we investigated physiological responses of 4-years-old *Tridacna maxima* specimens to temperature and *p*CO_2_ conditions in the French Polynesian lagoons during the today warmer season and those predicted for 2100 by the IPCC 2014: +1.5°C (RCP 4.5 scenario) and +800 *µ*atm of CO_2_ (RCP 8.5 scenario). Effect of each parameter and their potential synergetic effect were evaluated on respiration, biomineralization, and photophysiology by analyzing the holobiont O_2_ production and respiration, growth rate and ultrastructure of the shell and symbiont density and photosynthetic yield. In addition, the kinetics of thermal and acidification stress were accessed by performing analyses at different time of exposure (i.e., 29, 41, 53, 65 days).

## 2. Materials and Methods

### 2.1. Biological material

*Tridacna maxima* specimens of 4-years-old, 5-6 cm height and brownish/dark-green color, were collected in early January 2016 from the cultivated stock in Reao lagoon (Tuamotu islands) and exported to the Centre Ifremer du Pacifique in Tahiti where they were acclimatized in outdoor tank with running seawater. Each individual was placed onto a petri dish on which byssal gland further developed for fixation. All individuals directly opened up right after their transfer into the outdoor tank. No visual sign of stress was observed during the acclimation period except for 4 individuals which died within the first 3 days (corresponding to a 2% mortality rate). Specimens used in this study were collected and held under a special permit (MEI #284) delivered by the French Polynesian government.

### 2.2. Experimental design and rearing system

To study the impact of temperature, pCO_2_, and their putative synergetic effect on the physiology of the giant clams and their symbionts, four experimental conditions were set up by applying two temperatures (29.2°C and 30.7°C) and two levels of *p*CO_2_ (430 ±22 *μ*atm and 1212 ±35 *μ*atm). The tested conditions were: (1) control: 29.2°C 430 *μ*atm, (2) acidification stress: 29.2°C 1212 *μ*atm), (3) thermal stress: 30.7°C 430 *μ*atm, (4) acidification and thermal stress: 30.7°C 1212 *μ*atm (Table 1).

**Table 1.**
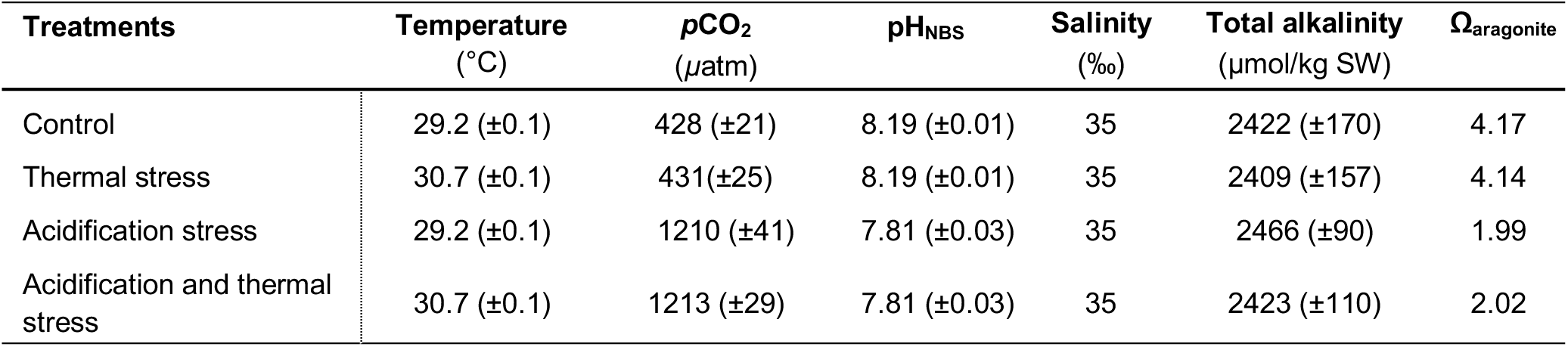
Measured and calculated parameters of seawater for all treatments. Total alkalinity is given in mean (±s.d.) based on weekly measurements for each experimental tank corresponding to n=36 for each condition. *p*CO_2_ and Ω_aragonite_ were calculated using the CO_2_ Systat software.

| Treatments | Temperature<br>(°C) | $p\text{CO}_2$<br>($\mu\text{atm}$) | $\text{pH}_{\text{NBS}}$ | Salinity<br>(‰) | Total alkalinity<br>($\mu\text{mol/kg SW}$) | $\Omega_{\text{aragonite}}$ |
| --- | --- | --- | --- | --- | --- | --- |
| Control | 29.2 ($\pm 0.1$) | 428 ($\pm 21$) | 8.19 ($\pm 0.01$) | 35 | 2422 ($\pm 170$) | 4.17 |
| Thermal stress | 30.7 ($\pm 0.1$) | 431 ($\pm 25$) | 8.19 ($\pm 0.01$) | 35 | 2409 ($\pm 157$) | 4.14 |
| Acidification stress | 29.2 ($\pm 0.1$) | 1210 ($\pm 41$) | 7.81 ($\pm 0.03$) | 35 | 2466 ($\pm 90$) | 1.99 |
| Acidification and thermal stress | 30.7 ($\pm 0.1$) | 1213 ($\pm 29$) | 7.81 ($\pm 0.03$) | 35 | 2423 ($\pm 110$) | 2.02 |

After a 3 weeks acclimation period in an outdoor tank, 96 clams were randomly distributed in experimental tanks one week before starting the experiment. For each condition, we used a 500 L tank containing 4 tanks of 30 L (ecological replicates) renewed at a flow rate of 50 L/h (seawater pumped in the lagoon). Each 30 L tank contained 6 clams (biological replicates). To avoid physiological shock, targeted temperature and *p*CO_2_ values were linearly achieved over 7 days. The light was set to obtain a Photosynthetically Active Radiation (PAR) of 200 ± 20 μmol of photons. m^-2^.s^-1^ on a 12 h:12 h light/dark photoperiod. To evaluate the kinetics of the thermal and/or acidification stress, analyses were performed at 4 different times of exposure, i.e., 29, 41, 53 and 65 days. In total, 64 clams were used for data acquisition corresponding to 4 individuals per condition (1 individual per 30 L tank) and per time of exposure.

### 2.3. Monitoring of temperature, pH and water quality

To insure the stability of experimental conditions, temperature and pH parameters were measured twice a day for each tank at 8:00 am and 4:00 pm using a mercury thermometer (certified ISO 9001, ±0.1°C accuracy) and a pH-meter Consort P603 (±0.01 accuracy). Total alkalinity (TA) was weekly titrated using a 0.01 N HCl solution and a titrator (Schott Titroline Easy). Levels of *p*CO_2_ and aragonite saturation state were calculated from temperature, pH (NBS scale), salinity and mean TA using the CO_2_ Systat software (van Heuven et al., 2009). All parameters including seawater carbonate chemistry are reported in Table 1.

### 2.4. Holobiont O_2_ consumption and production measurements

Giant clams were placed in an ecophysiological measurement system (EMS) to monitor O_2_ consumption and production. The EMS consisted in five open-flow chambers. Four giant clams were individually placed into 4 chambers while an empty shell was placed into a fifth chamber used as a control. EMS chambers contained water at the same temperature and *p*CO_2_ conditions as in the experimental tanks. The light energy and photoperiod conditions were the same as for the acclimation tanks. Flow rate in all chambers was constantly maintained at 12 L.h^-1^. Each chamber was equipped with a two-way electromagnetic valve activated by an automaton (FieldPoint National Instruments). When the electro-valve was opened, the water released from the chamber was analyzed for 3 min using an oxygen sensor (OXI 538, Cellox 325, WTW, Weilheim, Germany) to quantify dissolved oxygen. Oxygen measurements were performed over 48 h. The first 8 hours of measurement were discarded due to the animal acclimatization to the chamber. In each chamber, the cycle was completed within 3 min: the first 2 min served to stabilize the measurement and an average of oxygen data was performed on the last minute of acquisition. This cycle was followed by another time frame of 3 min in the control chamber following the sequence: specimen #1, control, specimen #2, control, specimen #3, control, specimen #4, control. Respiration rate (RR) and production rate (PR) were calculated from data obtained during night- and day-time, respectively, using differences in oxygen concentrations between the control and experimental chambers. RR and PR = V(O1-O2), with O1 the oxygen concentration in the control chamber, O2 the oxygen concentration in the experimental chamber, and V the water flow rate. RR and PR data were normalized to tissue dry weight. Once normalized, the terminology becomes O_2_ consumption for RR and O_2_ production for PR, both expressed in mg O_2_.h^-1^.g^-1^ dry weight.

After O_2_ production and O_2_ consumption analyses were completed, a piece of the mantle was dissected for further symbiont fluorescence and density analyses (see section 2.5). The remaining soft tissues were frozen and lyophilized for RR and PR data normalization.

### 2.5. Fluorescence and density measurements of symbionts

Potential effect of temperature and *p*CO_2_ conditions on the photophysiology of the symbionts was studied by comparing fluorescence yield of photosystem II between all experimental conditions. After clams were sacrificed, a 1 cm × 2 cm mantle fragment was dissected. The tissue fragment was gently swiped using tissue paper to remove a maximum of mucus and symbionts were collected by doing 5 smears using a sterilized razor blade. Collected symbionts were diluted into 5 mL of 0.2 µm filtered-seawater and placed at the obscurity for 10 min to inactive the photosystem II before light excitation. Samples were homogenized and 3 ml of the homogenate were collected, placed into a quartz-glass cuvette and analyzed with AquaPen fluorometer (APC-100, Photon System Instruments^®^, Czech Republic) at a 450 nm wavelength. The fluorescence at the steady state (F_0_) and the maximal fluorescence in the light (F_m_) were measured. The apparent quantum yield of photosynthesis (Y) reflecting the efficiency of photosystem II in the light acclimated state (Hoogenboom et al., 2006) was calculated according to equation (Eqn 1).

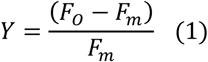

In addition, symbiont densities were evaluated from mantle fragments. For each individual, a circular (5 mm diameter) piece of mantle was collected using a punch. The piece was weighted, grounded in 0.2 µm filtered seawater and homogenized. Then, 20 µL of the tissue extract were immediately collected and placed into Mallassez cells for symbiont counting under optical microscope. For each sample, counting was performed on four replicates (3 columns per replicate). Data are expressed in number of symbionts/mg of mantle tissue.

### 2.6. Calcein labelling for evaluating daily shell extension rate

To study the impact of temperature and *p*CO_2_ on shell growth rate, the mineralization front of giant clams was marked using calcein fluorochrome which is irreversibly precipitated at the CaCO_3_ mineralization site. Before the experiment starts, giant clams were immersed in a 100 mg/L calcein solution (Sigma Aldrich) (calcein diluted in 1 µm filtered-seawater) for 8 h in the dark. During the labelling procedure, the bath of calcein solution was aerated using bubblers and a water current was created via pumps. Calcein-labelled specimens were then placed into the experimental tanks. At the end of the experiment, for each individual, a 5 mm thick section was cut along the maximal shell growth axis through the right valve using a Swap Top Inland^®^ diamond saw. All sections obtained were polished and observed under epifluorescence (λ_excitation_ = 495 nm, λ_emission_ = 515 nm) with a Leitz Dialux^®^ 22 microscope. The distance between the fluorescent calcein mark and the edge of the shell formed during the experiment was measured following the maximal growth direction. Daily shell extension rate (expressed in μm/day) was obtained by dividing the measured distance by the number of days of incubation in experimental conditions.

### 2.7. Shell scanning electron microscopy study

To characterize temperature and *p*CO_2_ effect on the shell ultrastructure, a scanning electron microscopy (SEM) study was carried out. For each individual, a 10 mm thick shell section was cut facing the section used for calcein observations. The section was then fractured along the width, 2 cm below the growing part of the shell (marginal part), using a small chisel and a hammer. Then, the apical fragment was longitudinally fractured and one piece was sonicated in tap water for 10 s, air-dried and an additional drying was done overnight at 35°C. Sample was placed on a stub covered with carbon tape, gold-coated and observed at 15 kV using a Hitachi TM3030 SEM at the Université de la Polynésie française. For each condition, and for each time of exposure, 2 samples were selected based on their daily extension rate, i.e., samples showing the lowest and the highest rate. To evaluate the impact of temperature and/or *p*CO_2_ on the shell ultrastructure, SEM observations were performed for each specimen in two different zones of the crossed-lamellar outer layer (Pätzold et al., 1991). Zone 1 corresponds to the shell formed *in situ* (i.e., in the shell region located before the calcein mark) while zone 2 corresponds to the shell formed during the experiment. Observations were made at the ultrastructural level and focused on the aspect and integrity of the lamellae of the crossed-lamellar outer layer.

### 2.8. Statistical analyses and data processing

Normality of data distribution and homogeneity of variance were tested with the Shapiro-Wilk test and the Bartlett test, respectively. Production and consumption of O_2_ data followed the conditions of application of parametric tests, but photosynthetic yields and symbiont densities were transformed using Box Cox transformation while shell extension rates were square root to meet these conditions. Comparisons were done using a three-way ANOVA with interaction where factors were time of exposure, temperature and *p*CO_2_. Tukey post hoc comparisons were done at α=0.05 for all analyses. Correlations between physiological parameters were tested using Pearson method with a threshold of r=0.25 (α=0.05).

For all physiological parameters, i.e., O_2_ respiration, O_2_ consumption, symbiont photosynthetic yield and density, means (±s.d.) were calculated based on the 4 biological replicates for each condition and each time of exposure. For the daily shell extension rate, means (±s.d.) were calculated for each condition and each time of exposure.

## 3. Results

### 3.1. Effect of temperature and pCO_2_ on the holobiont oxygen balance

For all sampling time (i.e., 29, 41, 53, 65 days), a cyclic pattern of O_2_ production and consumption is observed following the circadian cycle (Fig. 1). This pattern corresponds to oxygen photosynthetically produced and heterotrophically consumed by the holobiont during the day and the night, respectively. Mean values of normalized O_2_ production and consumption are shown in Fig. 2. These values were analyzed using a three-way ANOVA with 3 fixed factors: temperature, *p*CO_2_, and the time of exposure (Table 2). Tukey post hoc test results are reported in Table 3. The ANOVA indicates that oxygen production of the holobiont during day-time is significantly altered at 30.7°C (*P*=0.009) but not at 1212 *μ*atm of CO_2_ (*P*=0.361) (Table 2). In addition, O_2_ production is higher at 29.2°C than at 30.7°C for all times of exposure (Fig. 2A) and decreases along the experiment (*P*=0.030) (Table 2). The night-time O_2_ consumption, however, is not significantly influenced by temperature (*P*=0.590) neither by *p*CO_2_ (*P*=0.361) nor by time of exposure (*P*=0.533) and remains stable along the whole experiment (Fig. 2A,B). No interaction effect between the 3 tested factors is found to affect the holobiont oxygen balance (Table 2).

**Table 2.** Results from the three-way ANOVA performed on holobiont O_2_ production and consumption data, symbiont photosynthetic yield, symbiont density, and giant clam shell extension rate. The three fixed factors are *p*CO_2_, temperature and time of exposure. Significant differences (*P*<0.05) are highlighted in grey.

|  |  | O<sub>2</sub><br>production<br>(n=64) | O<sub>2</sub><br>consumption<br>(n=64) | Photosynthetic<br>yield<br>(n=64) | Symbiont<br>density<br>(n=64) | Shell extension<br>rate<br>(n=55) |
| --- | --- | --- | --- | --- | --- | --- |
| **Time** | *F* | 3.264 | 0.740 | 3.566 | 6.035 | 4.695 |
|  | *P* | 0.030 | 0.533 | 0.021 | 0.001 | 0.007 |
| **Temperature** | *F* | 7.551 | 0.294 | 65.200 | 79.158 | 0.466 |
|  | *P* | 0.009 | 0.590 | <0.0001 | <0.0001 | 0.499 |
| **pCO<sub>2</sub>** | *F* | 0.846 | 0.851 | 6.125 | 8.665 | 7.409 |
|  | *P* | 0.362 | 0.361 | 0.017 | 0.005 | 0.010 |
| **Time x Temperature** | *F* | 0.413 | 0.965 | 3.393 | 1.528 | 0.319 |
|  | *P* | 0.744 | 0.417 | 0.025 | 0.219 | 0.812 |
| **Time x pCO<sub>2</sub>** | *F* | 1.765 | 0.476 | 0.024 | 1.033 | 0.251 |
|  | *P* | 0.167 | 0.700 | 0.995 | 0.386 | 0.860 |
| **Temperature x pCO<sub>2</sub>** | *F* | 1.334 | 1.029 | 0.100 | 1.989 | 0.026 |
|  | *P* | 0.254 | 0.315 | 0.753 | 0.165 | 0.872 |
| **Time x Temperature x pCO<sub>2</sub>** | *F* | 1.216 | 0.148 | 2.153 | 0.474 | 0.336 |
|  | *P* | 0.314 | 0.930 | 0.106 | 0.702 | 0.799 |

**Table 3.**
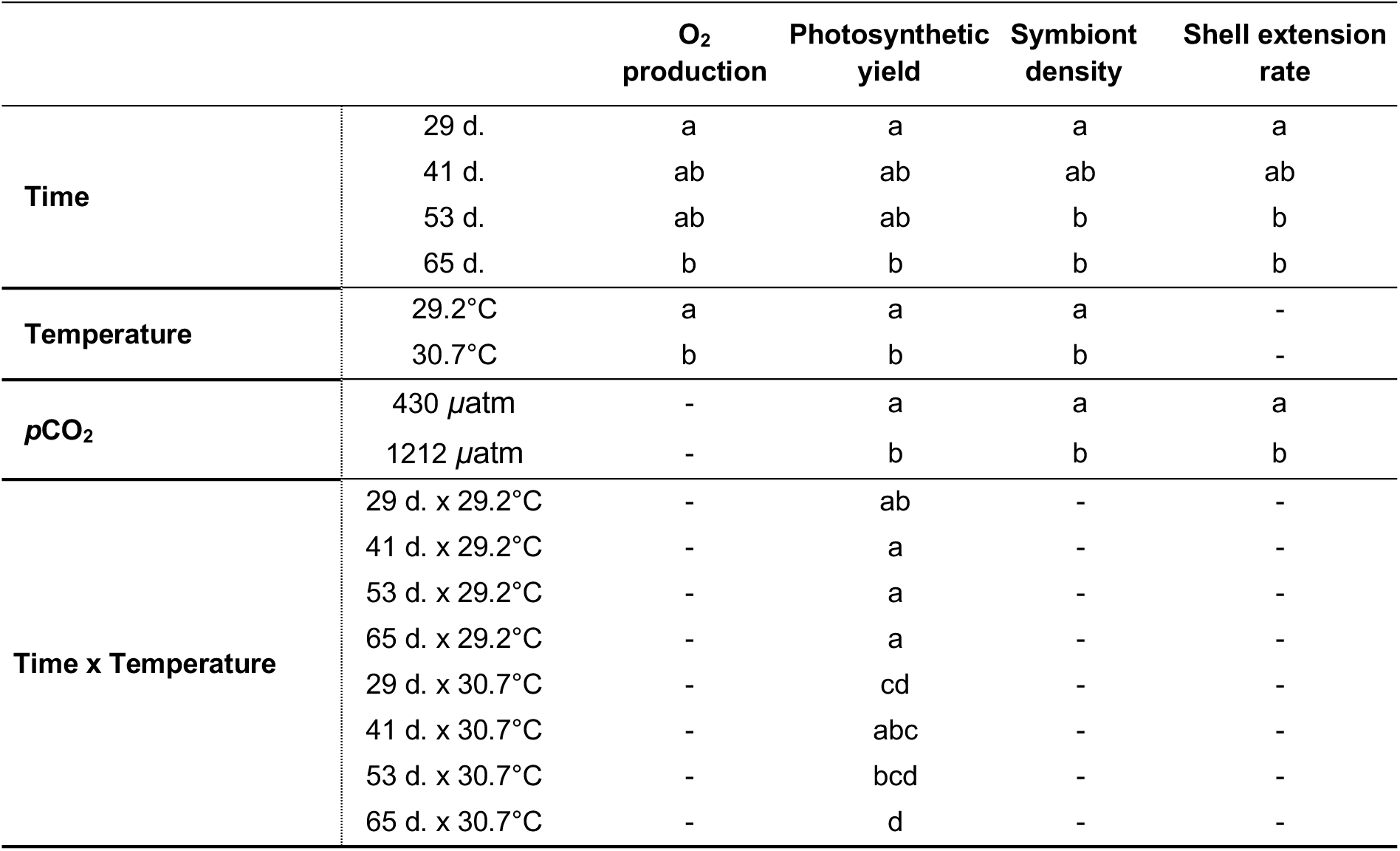
Results from Tukey post hoc tests following the three-way ANOVA performed on analyzed physiological parameters. The effects of significant parameters were tested as time of exposure alone and combined to the temperature. The letter annotations correspond to the significances between conditions (*P*<0.005).

**Figure 1.**
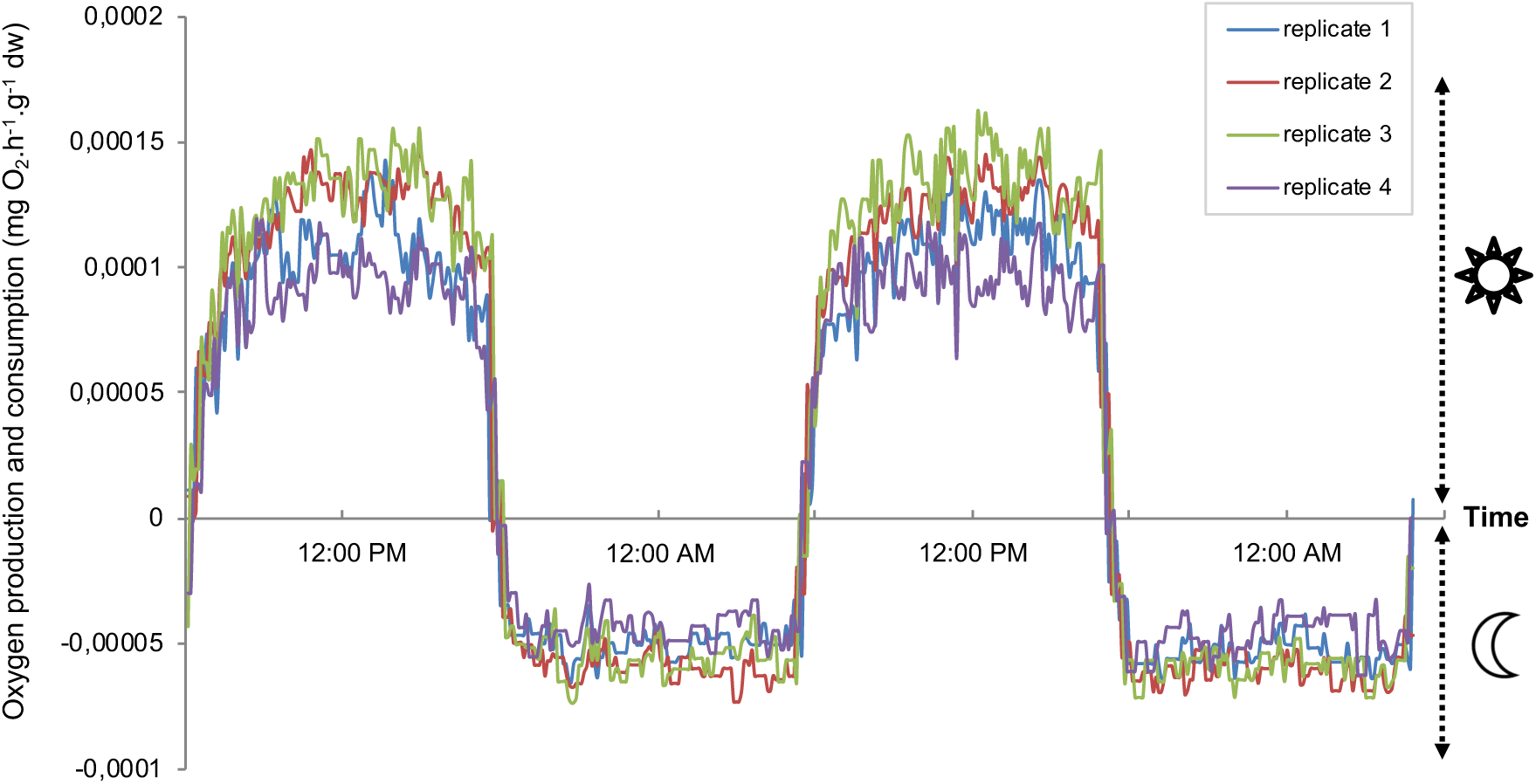
Variations of oxygen consumption and oxygen production acquired over a 48 h period after 29 days of exposure to 29.2°C and 430 *µ*atm of CO_2_. Data are expressed in mg O_2_.h^-1^.g^-1^ tissue dry weight (dw) and correspond to day-time and night-time acquisitions for 4 giant clam replicates.

**Figure 2.**
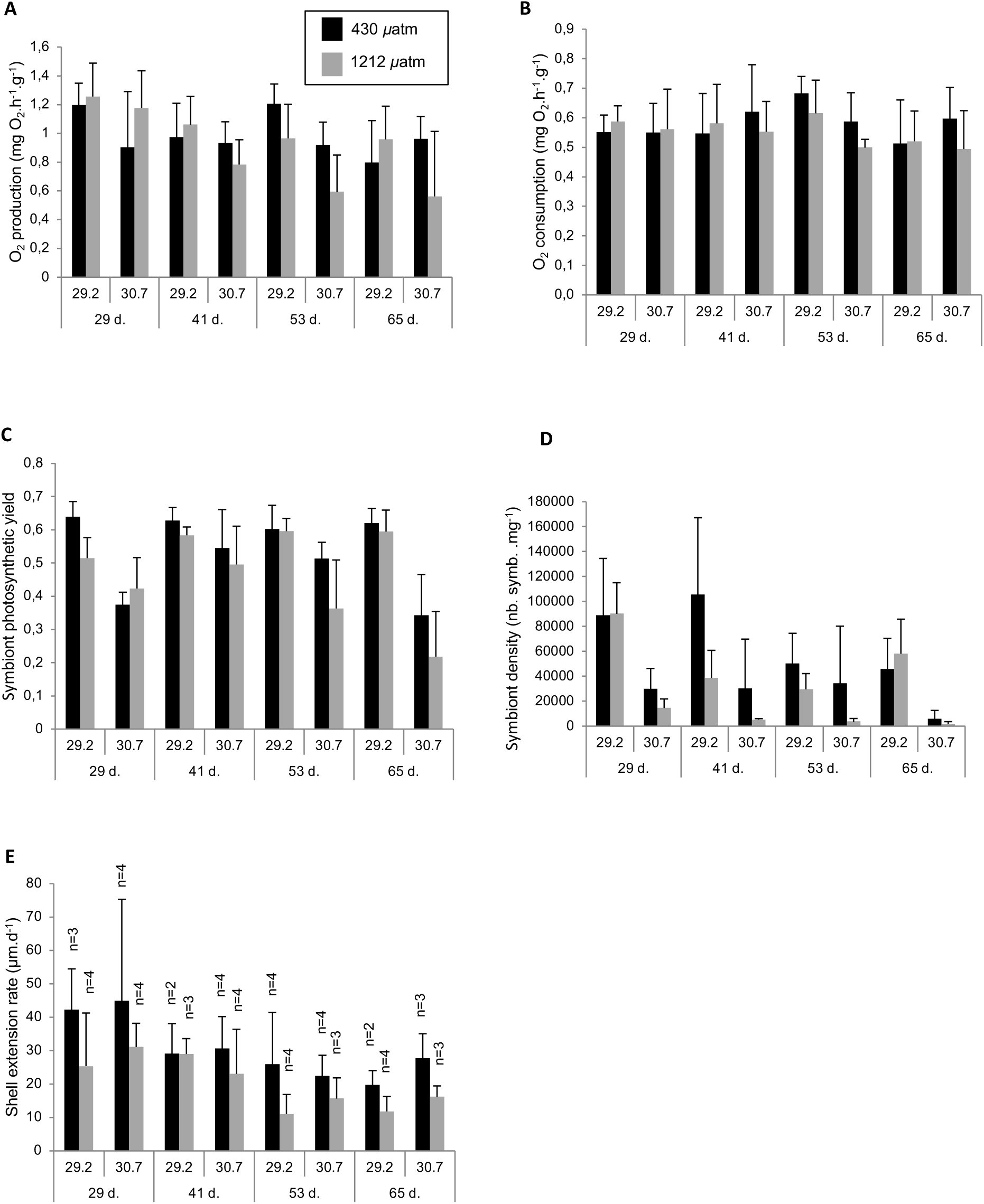
Graphs reporting data of (A) oxygen production, (B) oxygen consumption, (C) symbiont photosynthethic yield and (D) density, and (E) daily shell extension rate obtained for each temperature/*p*CO_2_ experimental condition and time of exposure. Black and grey columns correspond to today *p*CO_2_ condition (430 *µ*atm) and future *p*CO_2_ condition (1212 *µ*atm), respectively. Data are given in mean (+s.d.) calculated from 4 replicates (n=4) for each physiological parameter excepted for daily shell extension rate. For this latter parameter, numbers of replicates are indicated on the graph.

### 3.2. Symbiont density and photosynthesis under different T°/pCO_2_ conditions

After two months of exposure, the symbiont photosynthetic yield and density are significantly impacted at 30.7°C (*P*<0.0001), high *p*CO_2_ (1212 *μ*atm) (*P*<0.0001 and *P*=0.05, respectively), and by the time of exposure (*P*=0.021 and *P*=0.001, respectively). During the whole experiment and for both *p*CO_2_ conditions, the means of photosynthetic yield (Fig. 2C) and symbiont density (Fig. 2D) tend to be higher at 29.2°C than at 30.7°C. However, an inter-individual variability is observed.

Concerning the interaction parameters, only interaction between temperature and time of exposure parameters has a significant effect on the symbiont photosynthetic yield (*P*=0.025) (Table 2). No parameter interaction is found to affect the symbiont density (Table 2).

### 3.3. Negative effect of pCO_2_ on daily shell extension rate

Calcein mark was detectable in 55 over 64 shell sections. Statistical analyses on shell extension rate data indicate that *p*CO_2_ and the time of exposure have a significant effect on the shell extension rate (*P*=0.010 and *P*=0.007, respectively) (Table 2). In the Fig. 2E, the mean values of shell extension rates tend to be lower at high *p*CO_2_.

### 3.4. Relationship between physiological parameters

To establish if relationships exist between the various physiological parameters measured, a correlation matrix was generated (Table 4). The photosynthetic yield of symbionts is strongly correlated to their density (r=0.667). O_2_ production is also strongly correlated to symbiont density (r=0.482) and photosynthetic yield (r=0.331). Concerning the circadian functioning of the holobiont, its nocturnal oxygen need is strongly correlated to the diurnal O_2_ production (r=0.629). No significant correlation is found between shell extension rate and other physiological parameters.

**Table 4.**
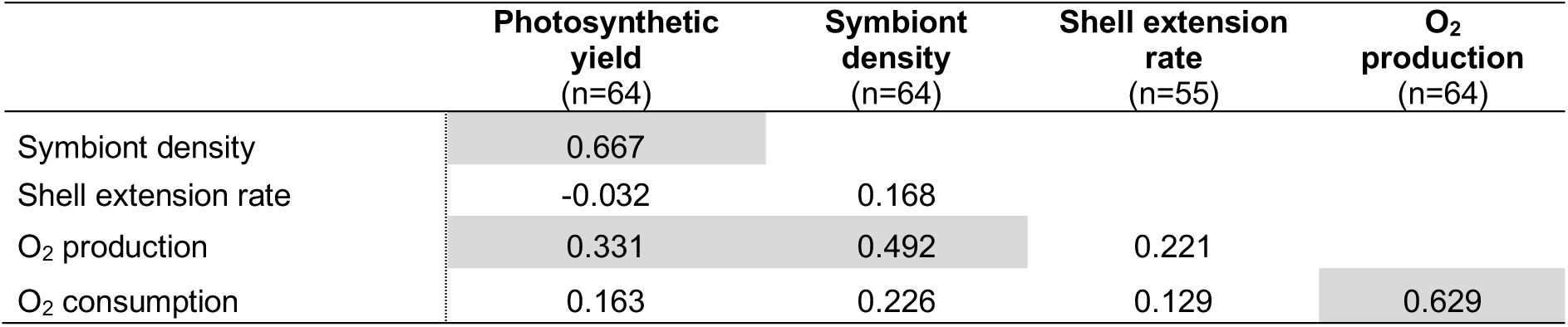
Correlation matrix integrating the different physiological parameters monitored, i.e., O_2_ production and consumption, shell extension rate and symbiont photosynthetic yield and density. Correlation were tested using the Pearson method (threshold of r=0.250, α=0.05) and significant relationships between parameters are highlighted in grey.

### 3.5. Effect of long-term exposure to temperature and pCO_2_ on the shell microstructure

Ultrastructural observations of shell fractures were done in two zones: zone 1 corresponding to the shell formed *in situ* (i.e., before the experiment) and zone 2 corresponding to the shell formed in the experimental conditions. The lamellae formed before the experiment (zone 1) are well-cohesive and display an elongated shape, a smooth surface, and sharp or slightly rounded outlines (Fig. 3A,B,C). For all experimental conditions, no difference between zone 1 and 2 is observed for all samples exposed for 29 days (Fig. 3A,D). From 41 days, the lamellae observed in the majority of shells formed during the experiment (zone 2), in all temperature/*p*CO_2_ conditions, appeared less cohesive with a pronounced granular aspect of the surface (Fig. 3E,H) and/or less cohesive lamellae with a crazed-like aspect (Fig. 3F,I). Two states classifying ultrastructural differences between the shell formed before and during the experiment were defined as follow: *i*) ND; no difference observed between zone 1 and 2, *ii*) D; difference observed between zone 1 and 2. In total, 19 samples over 24 displayed differences in lamellae aspect between both zones. Data are reported in Table 5.

**Table 5.** Compilation of ultrastructural observations of the lamellae of the shell outer layer using scanning electron microscopy. “ND” means that no significant difference is observed between the shell formed before and during the experiment. “D” means that a notable difference is observed between both parts of the shell. “A” and “B” represent the two samples observed for each treatment and time of exposure.

| Time of exposure | Temperature (°C) | pCO<sub>2</sub> (μatm) | Sample | Difference |
| --- | --- | --- | --- | --- |
| 29 d. | 29.2 | 430 | A | ND |
|  |  |  | B | ND |
|  | 30.7 | 430 | A | ND |
|  |  |  | B | ND |
|  | 29.2 | 1212 | A | ND |
|  |  |  | B | ND |
|  | 30.7 | 1212 | A | ND |
|  |  |  | B | ND |
| 41 d. | 29.2 | 430 | A | D |
|  |  |  | B | D |
|  | 30.7 | 430 | A | D |
|  |  |  | B | D |
|  | 29.2 | 1212 | A | D |
|  |  |  | B | D |
|  | 30.7 | 1212 | A | D |
|  |  |  | B | ND |
| 53 d. | 29.2 | 430 | A | D |
|  |  |  | B | D |
|  | 30.7 | 430 | A | D |
|  |  |  | B | D |
|  | 29.2 | 1212 | A | ND |
|  |  |  | B | D |
|  | 30.7 | 1212 | A | D |
|  |  |  | B | ND |
| 65 d. | 29.2 | 430 | A | D |
|  |  |  | B | D |
|  | 30.7 | 430 | A | D |
|  |  |  | B | ND |
|  | 29.2 | 1212 | A | D |
|  |  |  | B | D |
|  | 30.7 | 1212 | A | ND |
|  |  |  | B | D |

**Figure 3.**
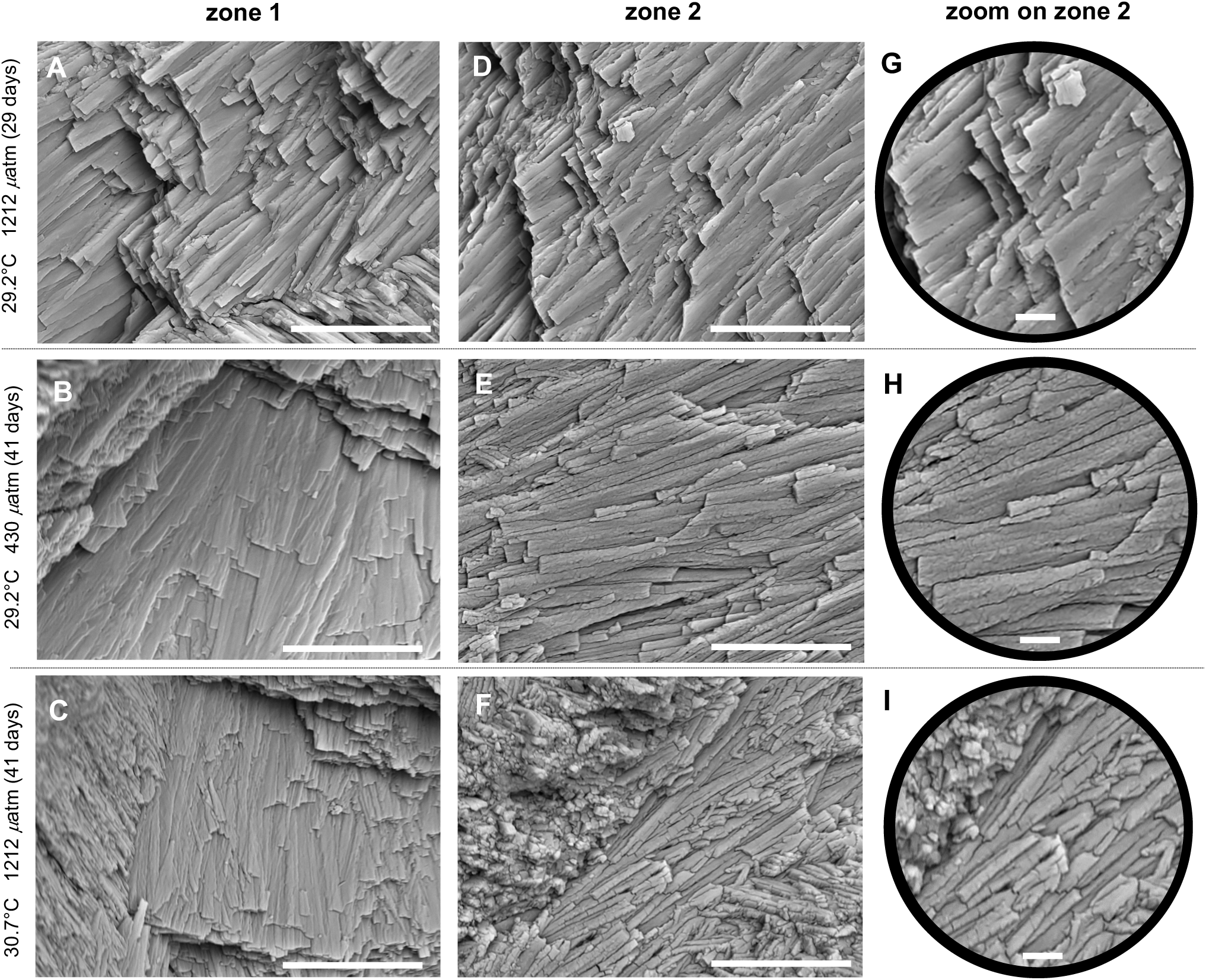
Effects of experimental conditions on the ultrastructure of the *Tridacna maxima* shell outer layer investigated via scanning electron microscopy. (A-F: scale bar: 10 µm, G-I: scale bar: 2 µm). Zone 1 and zone 2 correspond to the shell formed *in situ* (i.e., before the experiment) and in the experimental conditions, respectively. (A, B and G): 29.2°C 1212 *µ*atm (29 days): no difference between zone 1 and 2. In both zones, lamellae are well-cohesive, displaying elongated shape with a smooth surface and slightly rounded outlines. (B, E and H) 29.2°C 430 *µ*atm (41 days): lamellae in zone 2 are less-cohesive, with a pronounced granular aspect. (C, F and I) 30.7°C 430 *µ*atm (41 days): lamellae in zone 2 are less-cohesive and display crazes.

### 3.6. Effect of time of exposure on oxygen balance, biomineralization and symbionts photophysiology

Time of exposure to thermal and acidification stress was shown to have a significant impact on O_2_ production (*P*=0.030), photosynthetic yield (*P*=0.021), symbiont density (*P*=0.001) and shell growth rate (*P*=0.007) (Table 2). Post hoc tests showed that the kinetics of thermal or acidification stress varies depending on the physiological parameter measured: O_2_ production and photosynthetic yield are significantly different between 29 and 65 days while symbiont density and shell growth rate are both impacted from 53 days of exposure (Table 3).

## 4. Discussion

In this present study, we investigated the physiological responses of *Tridacna maxima* (i.e., 4-years-old specimens) to temperature and *p*CO_2_ conditions in the French Polynesian lagoons under nowadays warmer season conditions and those predicted for 2100 by the IPCC 2014 (+1.5°C and +800 *µ*atm of CO_2_). Today thermal conditions are already and sufficiently stressful to induce an acclimatization process pictured by a slight regulation of the symbiont density without the collapse of the photosynthesis. However, if in the today conditions the giant clams are able to mount this acclimatory and efficient response that enable to maintain their key physiological characteristics, the thermal and *p*CO_2_ conditions of tomorrow appear to be much more stressful and sufficient to induce strong disturbances of the two key functional compartments of the giant clam, its phototrophic symbiosis and its biomineralization. Indeed, the 30.7°C treatments have significantly impacted the holobiont by inducing a strong bleaching response illustrated by the reduction of its O_2_ production, symbiont density and photosynthetic yield. High *p*CO_2_ (+800 *µ*atm of CO_2_) was shown to alter the symbiont photosynthetic yield and density and to affect its biomineralization process by decreasing the shell growth rate.

### 4.1. No synergetic effect between temperature and pCO_2_ on giant clam and symbiont physiological parameters

Increase of atmospheric CO_2_ impacts both the seawater temperature and pH inducing ocean warming and acidification (Sabine et al., 2004, IPCC, 2014). Therefore, the analysis of the synergetic effect of both stressors on marine bivalve physiology is crucial. Watson et al. (2012) investigated the synergetic effect of ocean acidification and warming on the survival of *T. squamosa* juveniles over a 60 days crossed-factor experiment. They reported that the combination of the highest *p*CO_2_ (+600 *µ*atm of CO_2_) and both moderate and highest seawater temperatures (30.0°C and 31.5°C) resulted in the lowest survival rate of giant clams (<20%). They concluded that increased ocean *p*CO_2_ and temperature are likely to reduce the survival of *T. squamosa* juveniles. In our study, no synergetic effect was detected for all measured physiological parameters on *T. maxima*. Moreover, no mortality was observed in all tested temperature/*p*CO_2_ conditions during the whole experiment which lead us to suggest that a temperature of 30.7°C and a *p*CO_2_ of 1212 *µ*atm seem to be non-lethal, at least over 65 days of exposure. However, we cannot exclude that some specimens, especially the totally bleached individuals, may have died after a longer exposure to the experimental conditions.

### 4.2. Effect of temperature on holobiont oxygen balance, symbiont photophysiology and shell ultrastructure

Temperature is a factor well-known to influence several physiological parameters in bivalves as it regulates the metabolism of ectotherm organisms (Aldridge et al., 1995, Bougrier et al., 1995, Watson et al., 2012, Gazeau et al., 2013; Le Moullac et al., 2016a; Latchère et al., 2017). However, since giant clams live in symbiosis with photosynthetic symbionts (Klumpp et al., 1992; Klumpp and Griffits, 1994; Soo and Todd, 2014), the comparison at the metabolic level with non-symbiotic bivalves can be audacious. Discussion on elevated temperature effect on the holobiont oxygen balance should therefore integrate the effect on both giant clam and its symbionts (Jones and Hoegh-Guldberg, 2001). Even though physiological and molecular response to thermal stress may differ, the comparison with scleractinian corals, which are also symbiotic and calcifying organisms, makes sense to better understand the impact of temperature on the holobiont physiology.

We observed that high temperature (+1.5°C) significantly reduces the holobiont O_2_ production, the density and the photosynthetic yield of symbionts from 29 days of exposure. Additionally, partial or total bleaching was observed for the majority of individuals exposed to 30.7°C (at both ambient and high *p*CO_2_, Fig. S2). This suggests that thermal stress has a significant impact on the holobiont photophysiology. The reduction of the symbiont photosynthetic yield could reflect a photoinhibition. Photoinhibition was previously linked to the degradation of the D1 protein of the reaction center of the photosystem II (PSII) altering the photosynthetic apparatus functioning in symbiotic corals and sea anemone subjected to thermal stress (Warner et al., 1999; Smith et al., 2005; Richier et al., 2006; Ferrier-Pagès et al., 2007). Diminution of photosynthetic activity reduces the symbiont O_2_ production and consequently, impacts the holobiont O_2_ production. Comparable results were obtained from a 24-hour experiment showing that an increase of 3°C affects the oxygen production of *Tridacna gigas* and *Tridacna deresa* (Blidberg et al., 2000). Photoinhibition led to the production of reactive oxygen species (ROS) by symbionts which are known to pass through cellular membranes, cause oxidative damages (Lesser et al., 1990; Lesser, 1996; Downs et al., 2002) and impact the photosystem II (Richter et al., 1990). Considering corals, a strong positive correlation between accumulation of oxidative damage and coral bleaching was shown (Downs et al., 2002). Indeed, ROS such as hydrogen peroxide (H_2_O_2_) may play a role of signaling molecule activating the symbiosis dissociation (Fitt et al., 2001; Smith et al., 2005; Perez and Weis, 2006). Expulsion of symbionts by the host may be a strategy to limit oxidative stress and damage to ultimately survive environmental stress (Downs et al., 2002; Perez and Weis, 2006). Two recent works support this hypothesis. One of Zhou et al. (2018) which related a decrease in symbiont density in response to an excess of oxidative stress in thermally stressed *T. crocea*. A second of Dubousquet et al. (2016) which showed that *T. maxima* overexpressed genes encoding ROS scavengers in response to thermal stress. Interestingly, the cellular and molecular mechanisms enhanced upstream the bleaching response seem similar between giant clams and corals and conduct to the same phenomenon of symbiosis dissociation. However, since giant clams and corals display an extra- and an intra-cellular symbiosis respectively, the mechanisms of the symbiosis dissociation must differ. In every instance, this constitutes an interesting example of evolutionary convergence of a stress response in two very distant organisms. In this context, it would be interesting to test if the bleaching response in giant clam can be adaptive as it was proposed for corals (Fautin and Buddemeier, 2004) and if variations in giant clam thermo-tolerance are correlated to the composition of the symbiotic population (Baker, 2003).

Concerning the effect of temperature on the ultrastructure of the crossed-lamellar structure, the lamellae formed in all experimental conditions in the first 29 days are well-cohesive with elongated shape and a granular aspect which is consistent with the description made by Agbaje et al. (2017) in *T. deresa*. However, after 29 days, the lamellae observed in the majority of shells formed in both temperature conditions (whatever the *p*CO_2_ condition) appeared to be less-cohesive with pronounced granular aspect. We suggest that these features are due to a lack of organic matrix between the lamellae (i.e., inter-lamellar organic matrix) and embedding the nano-grains forming the first-order lamellae in the crossed-lamellae structure (as described by Agbaje et al., 2017). Indeed, the biomineralization of calcium carbonate structures involved the transport of ions (calcium, bicarbonates and protons) to the mineralization site and the synthesis of macromolecules referred to as “organic matrix” (Allemand et al., 2004). The organic matrix, consisting of 0.9% weight of *T. deresa* shell, and is mainly composed of assemblage of macromolecules such as polysaccharides, glycosylated, and unglycosylated proteins and lipids (Agbaje et al., 2017). These macromolecules are commonly found in molluscan shell organic matrix (Marin et al., 2012) present as inter-crystalline envelope and around individual crystal units (Harper, 2000). The formation of these macromolecules is energetically costly for mollusks such as marine gastropods (Palmer, 1992). According to the conclusion made by Milano et al. (2016) in the non-symbiotic bivalve *Cerastoderma edule*, we suggest that in our tested experimental temperature, while respiration function is significantly altered, less energy is available and allocated to the synthesis of macromolecules involved in biomineralization which may explained the lack of organic matrix embedded into the giant clam shell.

### 4.3. Effect of pCO_2_ on giant clam biomineralization and symbiont photophysiology

Ocean acidification is related to an increase of dissolved CO_2_ in the seawater resulting in a modification of carbonate chemistry equilibrium and a decrease of seawater pH. This phenomenon is known to affect biomineralization process of marine calcifiers since the production of biocarbonate structures depends on both calcium carbonate saturation state and on the ability to express proteins involved in biomineralization (Fitzer et al., 2014). The *p*CO_2_ is a well-known parameter to influence mollusk calcification process in terms of growth rate, expression of gene encoding proteins involved in biomineralization, and to alter the integrity of biocarbonate structures (Welladsen et al., 2010; Melzner et al., 2011; Fitzer et al., 2014; Le Moullac et al., 2016b), even if counter examples exist (Ries et al., 2009; Kroeker et al., 2010).

Concerning giant clams, our results indicate that in high *p*CO_2_ condition (+800 *µ*atm), shell extension rate decreases significantly. This observation is in accordance with those obtained for giant clams in three different experiments using various CO_2_ enrichment level and exposure duration. Waters (2008) and Watson (2015) have exposed juveniles *T. squamosa* to +200 *µ*atm of CO_2_ for 13 weeks and to +250 and +550 *µ*atm of CO_2_ for 8 weeks, respectively. Toonen et al. (2012) demonstrated that young specimens of *T. maxima* and *T. squamosa* have lower shell growth rates, compared to those reported in the literature under natural *p*CO_2_/pH conditions, when kept 1 year in +350 to +1000 *µ*atm *p*CO_2_ conditions.

As mentioned above, high *p*CO_2_ condition (+800 *µ*atm) affects giant clam shell growth rate but also symbiont photosynthetic yield and density. Negative effect of *p*CO_2_ on the shell growth rate may be linked to (1) the aragonite saturation state which is lower at 1212 *µ*atm than at 430 *µ*atm and/or (2) an alteration of symbiont photophysiology leading to a potential reduction of the ‘light-enhanced calcification’ (LEC). Concerning the former, decline of coral calcification rate has been related to modifications of carbonate chemistry due to a limitation of CO_3_^2-^ ions available for calcification. Even if corals may upregulate the pH at their mineralization site (Venn et al., 2013; Holcomb et al., 2014), the decrease of coral calcification was also linked to a decline in pH in the calcifying fluid due to a lower seawater pH (Ries et al., 2011; McCulloch et al., 2012). In our case, we suggest that giant clam *T. maxima* exposed to high *p*CO_2_ may allocate more energy to maintain a proper pH of the fluid in the extra-pallial space for nucleation and deposition of aragonite since the Ω_aragonite_ at high *p*CO_2_ is twice lower than at ambient *p*CO_2_ (Ω_aragonite, 1212 *µ*atm_ = 2, Ω_aragonite, 430 *µ*atm_ = 4). Regarding the light-enhanced calcification (LEC) (Vandermeulen et al., 1972), this phenomenon observed in Symbiodiniaceae-host symbiosis has been extensively described in corals and represents the capacity of symbionts to stimulate the host calcification (Allemand et al., 2004). The symbionts stimulate the host metabolism and calcification by providing energy resources and/or O_2_ (Chalker and Taylor, 1975). Moreover, symbionts may promote the aragonite precipitation by providing inorganic carbon, nitrogen and phosphorus and by synthetizing molecules used as precursor for the synthesis of skeletal organic matrix (Pearse and Muscatine, 1971; Cuif et al., 1999; Furla et al., 2000; Muscatine et al., 2005). They also facilitate CaCO_3_ precipitation by influencing the Dissolved Inorganic Carbon (DIC) equilibrium by removing the CO_2_ via photosynthesis (Goreau, 1959). Such phenomenon has been also described in giant clams (Ip et al., 2006; Ip et al., 2017). In *T. squamosa*, LEC increases the pH and reduces the ammonia concentration at the interface between the inner mantle and the shell in the extra-pallial fluid, where the biomineralization occurs (Ip et al., 2006). Recently, Chew et al. (2019) reported a light-enhanced expression of a carbonic anhydrase (i.e., CA4-like) in the inner mantle of *T. squamosa* and suggested that this enzyme is involved in giant clam biomineralization by catalyzing the conversion of HCO ^-^ to CO_2_. In this context, an altered photophysiology of the symbiont can rationally alter LEC and consequently results in a decrease of the shell growth rate. Finally, one can suggest that under acidification stress, giant clam may reduce or stop some physiological functions such as biomineralization and allocate more energy to essential functions to its survival. In our study, temperature also altered photophysiology and holobiont O_2_ production, but did not significantly affect shell growth rate. Therefore, the most plausible hypothesis explaining the negative effect of *p*CO_2_ on shell growth rate may be related to the low aragonite saturation state at high *p*CO_2_.

Concerning the effect of *p*CO_2_ on the shell microstructural integrity, in the temperate bivalve *Mytilus edulis*, an exposition to +150, +350 and +600 *µ*atm of CO_2_ for 6 months induced a disorientation in the shell of the newly formed calcite crystals of the prismatic layer (Fitzer et al., 2014). In the giant clam *T. maxima*, high *p*CO_2_ had no negative effect on the integrity of the aragonitic lamellae of the crossed-lamellar layer during the first 29 days of exposure. From 41 days of exposure, its potential impact remains unresolved as differences were noticed in the shells formed under future high temperature, high *p*CO_2_ and even under today high temperature/*p*CO_2_ conditions. The fact that differences are reported for shells formed in all experimental conditions from 41 days suggests that they may be due to a long-term exposure to 29.2°C and 30.7°C.

This study enables the evaluation of *Tridacna maxima* physiological responses to temperature and *p*CO_2_ thresholds predicted at the end of this century. We demonstrated that high temperature has a significant negative impact on symbiont densities and photosynthetic capacities, which induce a decrease in holobiont O_2_ production rate. We suggest that the observed decrease of holobiont O_2_ production rate results from a decrease of symbiont O_2_ production. Therefore, by influencing symbiont physiology, the temperature may affect the energetic needs of the giant clam host. The high *p*CO_2_ has a negative impact on shell growth rate, symbiont densities and photosynthetic capacities. Shell microstructure is affected neither by temperature nor by the *p*CO_2_ in the first 29 days of exposure. However, for all temperature/*p*CO_2_ conditions, a longer exposition (≥41 days) modified the shell ultrastructure. These observations support our hypothesis that 29.2°C is a temperature that already affects giant clam metabolism, at least over a long-term exposition. However, no synergetic effect was found between temperature and *p*CO_2_ parameters. All these observations suggest that temperature and *p*CO_2_ influence different physiological functions and that giant clam populations may dramatically suffer from the temperature and *p*CO_2_ conditions predicted for the next decades. To complement these results on the effects of temperature on the holobiont, it is now essential to conduct further and integrative analyses. This may include the definition of thermal optimum of *T. maxima* and the study of the transcriptomic response of the giant clam and its symbiotic Symbiodiniaceae to thermal stress for instance. All these analyses will allow a better understanding of the fundamental physiological processes of the holobiont and its response to future changes. Attempted results may also help in adapting local policies and management to maintain a sustainable exploitation of giant clam resource, especially in the Eastern Tuamotu islands where bleaching events have been observed at an increasing and alarming rate.

## Acknowledgments

The authors would like to thank Celine Lafabrie for fruitful discussions and Mickael Mege and Alexia Pihier for their technical assistance. Georges Remoissenet (Direction of Marine Resources of French Polynesia) is thanked for sharing information about bleaching events in Eastern Tuamotu islands. The Direction of Marine Resources and the Ministry of Economic Recovery, Blue Economy and Digital Policy of French Polynesia are also thanked for according the special permit to collect and hold small specimens of *Tridacna maxima* species. This study is set within the framework of the “Laboratoires d’Excellence (LabEX)” TULIP (ANR-10-LABX-41).

## Supplementary Material

**Figure S1.**
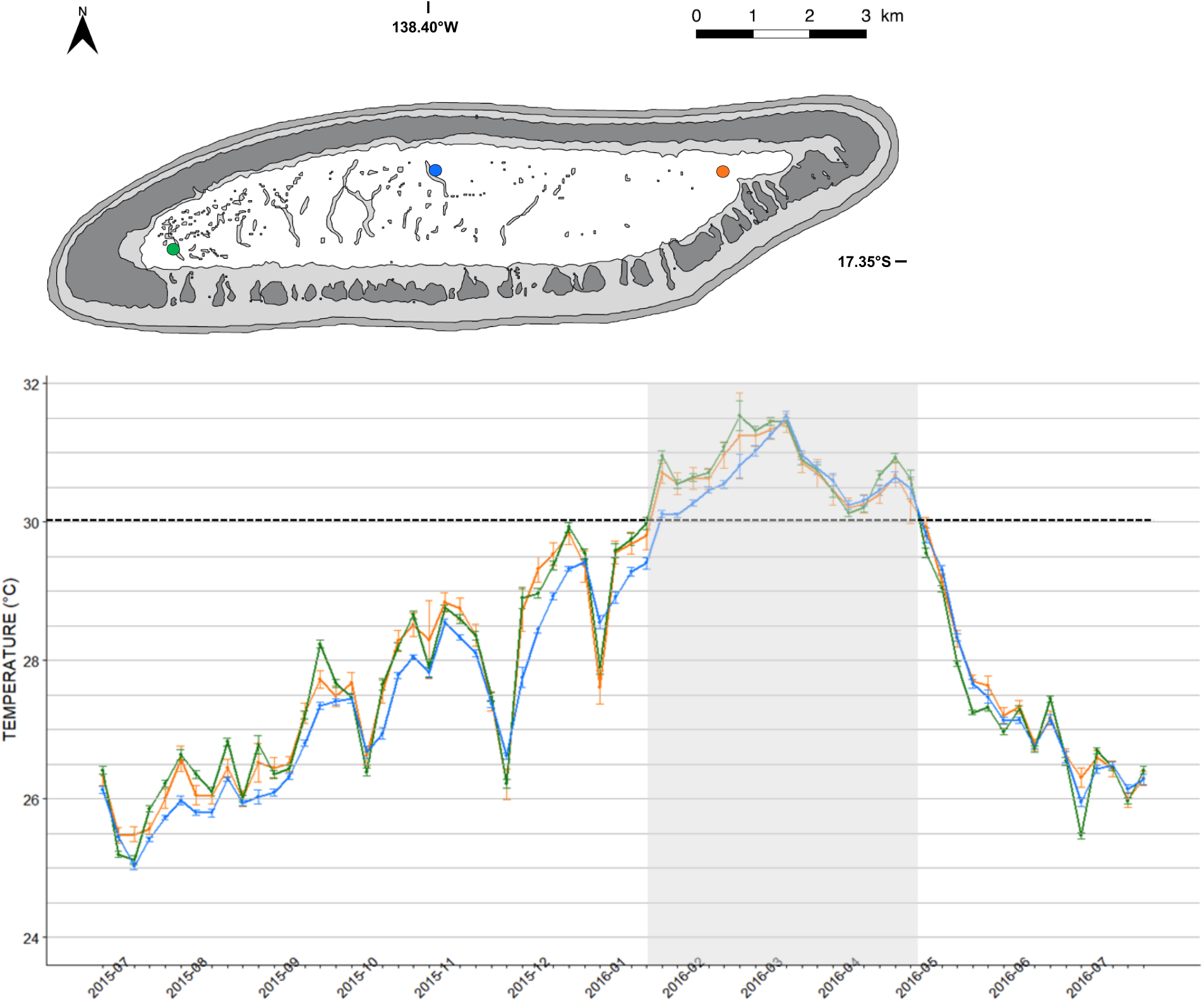
*In situ* seawater temperatures recorded from July 2015 to July 2016 inside the lagoon of Tatakoto atoll (Tuamotu islands) at 2 m depth. Temperature were recorded using UA-002-64 HOBO Onset data loggers. Curves on the graph report data from 3 sites localized on the map by circles with corresponding color. Grey rectangle highlights the period when giant clams were exposed to a temperature ≥30°C.

**Figure S2.**
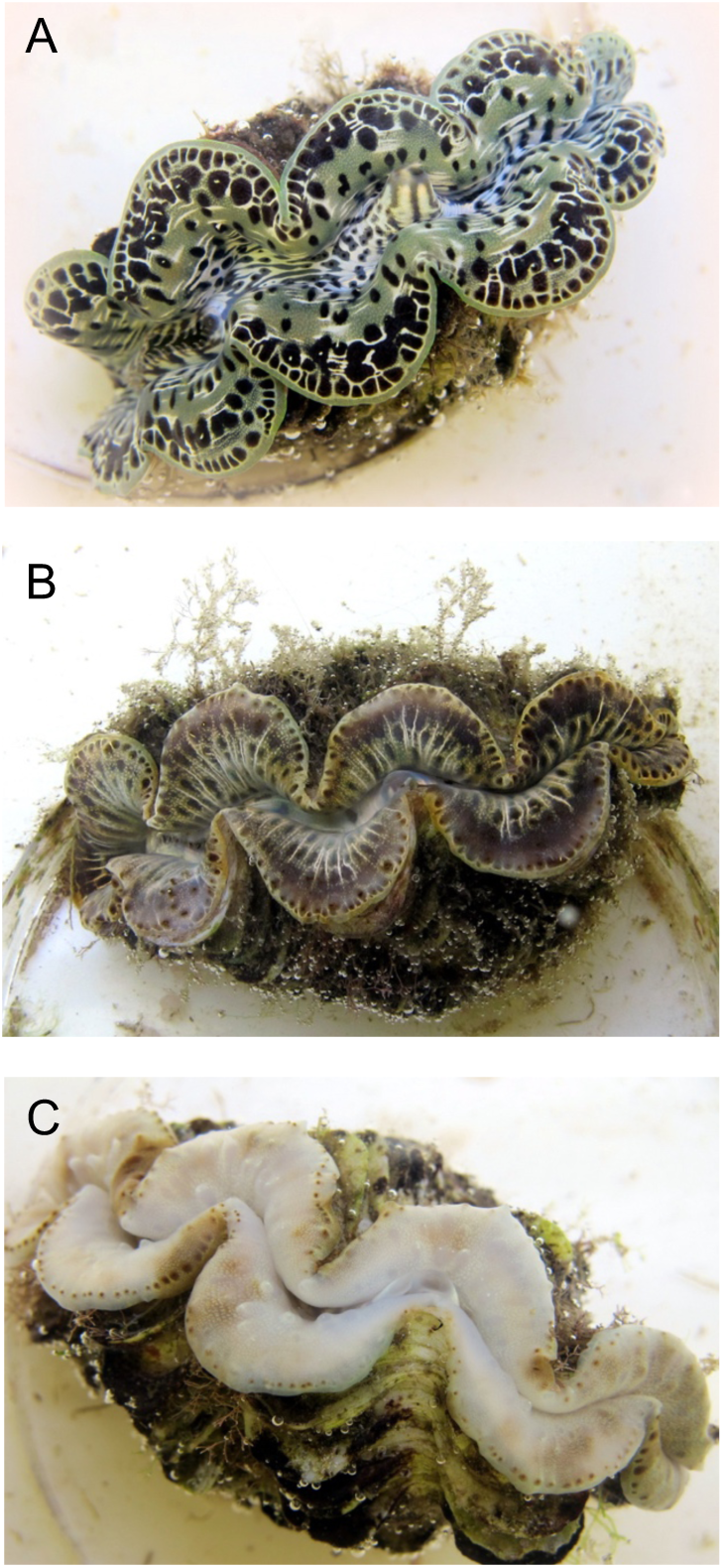
Visual observation of bleaching in giant clam *Tridacna maxima* specimens after 65 days in experimental conditions. (A) Non-bleached specimen exposed to 29.2°C 430 *µ*atm of CO_2_, (B) partially bleached and (C) completely bleached specimens exposed to 30.7°C 1212 *µ*atm of CO_2_.

## References

Adessi, L. (2001). Giant clam bleaching in the lagoon of Takapoto atoll (French Polynesia). Coral Reefs 19, 220.

Agbaje, O.B.A., Wirth, R., Morales, L.F.G., Shirai, K., Kosnik, M., Watanabe, T. and Jacob, D.E. (2017). Architecture of crossed-lamellar bivalve shells: the southern giant clam (*Tridacna deresa*, Röding, 1798). R. Soc. Open Sci. 4, 170622

Allemand, D., Ferrier-Pagès, C., Furla, P., Houlbreque, F., Puverel, S., Reynaud, S. Tambutté, E., Tambutté, S and Zoccola, D. (2004). Biomineralisation in reef-building corals: from molecular mechanisms to environmental control. C.R. Palevol. 3, 453–467.

Alcazat, S.N. (1986). Observations on predators of giant clams (Bivalvia: family Tridacnidae). Silliman J. 33, 54–57.

Aldridge, D.W., Payne, B.S. and Miller, A.C. (1995). Oxygen-consumption, nitrogenous excretion, and filtration-rates of *Dreissena polymorpha* at acclimation temperatures between 20 and 32 degrees. Can. J. Fish. Aquat. Sci. 52, 1761–1767.

Amstrong, E. (2017) Ion-regulatory and developmental physiology of giant clams (Genus Tridacna) and their conservation status on the island of Mo’orea, French Polynesia. PhD thesis, University of Berkley, USA.

Andréfouët, S., Van Wynsberge, S., Gaertner-Mazouni, N., Menkes, C., Gilbert, A. and Remoissenet, G. (2013). Climate variability and massive mortalities challenge giant clam conservation and management efforts in French Polynesia atolls. Biol. Conserv. 160, 190–199.

Andréfouët, S., Dutheil, C., Menkes, C.E., Bador, M. and Lengaigne, M. (2015). Mass mortality events in atoll lagoons: environmental control and increased future vulnerability. Glob. Chang. Biol. 21, 95–205.

Andréfouët, S., Van Wynsberge, S., Kabbadj, L., Wabnitz, C.C.C., Menkes, C., Tamata, T., Pahuatini, M., Tetairekie, I., Teaka, I., Ah Scha, T., Teaka, T. and Remoissenet, G. (2017). Adaptative management for the sustainable exploitation of lagoon resources in remote islands: lessons from a massive el Nino-induced giant clam bleaching event in the Tuamotu atolls (French Polynesia). Env. Conserv. 45, 30–40.

Baker, A. C. (2003). Flexibility and specifity in coral-algual symbiosis: Diversity, ecology, and biogeography of *Symbiodinium*. Annu. Rev. Ecol. Evol. Syst. 34, 661–689.

Bell, J.D., Lane, I., Gervis, M., Soule, S. and Tafea, H. (1997) Village-based farming of the giant clam *Tridacna gigas* (L.), for the aquarium market: Initial trials in Solomon Islands. Aquacult. Res. 28, 121–128.

Blidberg, E., Elfwing, T., Planhnan, P. and Tedengren, M. (2000). Water temperature influences on physiological behaviour in three species of giant clams (Tridacnidae). Proceeding 9th International Coral Reef Symposium, Bali, Indonesia, 1.

Broeker, W.S., Takahashi, T., Simpson, H.J. and Peng, T.-H. (1979). Fate of fossil fuel carbon dioxide and the global carbon budget. Science 206, 409–418.

Bougrier, S., Geairon, P., Deslouspaoli, J.M., Bacher, C. and Jonquieres, G. (1995). Allometric relationships and effects of temperature on clearance and oxygene consumption rates of *Crassostrea gigas* (Thunberg). Aquaculture 134, 143–154.

Buck, B.H., Rosenthal, H. and Saint-Paul, U. (2002). Effect of increased irradiance and thermal stress on the symbiosis of *Symbiodinium microadriaticum* and *Tridacna gigas*. Aquat. Living Resour. 15, 107–117.

Caldeira, K. and Wickett, M.E. (2003). Anthropogenic carbon and ocean pH. Nature 425, 365.

Chalker, B.E. and Taylor, D.L. (1975). Light-enhanced calcification, and the role of oxidative phosphorylation in calcification of the coral *Acropora cervicornis*. Proc. R. Soc. Lond. 190, 323–331.

Chew, S.F., Koh, C.Z.Y., Hiong, K.C., Choob, C.Y.L, Wong, W.P., Neo, M.L. and Ip, Y. (2019). Light-enhanced expression of Carbonic Anhydrase 4-like supports shell formation in the fluted giant clam *Tridacna squamosa*. Gene 683, 101–112.

Cuif, J.-P., Dauphin, Y., Freiwald, A., Gautret and P., Zibrowius, H. (1999). Biochemical markers of zooxanthellae symbiosis in soluble matrices of skeleton of 24 Scleractinia species. Comp. Biochem. Physiol. 123A, 269–278.

DeBoer, T.S., Baker, A.C., Erdmann, M.V., Jones, P.R., and Barber, P.H. (2012). Patterns of *Symbiodinium* distribution in three giant clam species across the biodiverse Bird’s Head region of Indonesia. Mar. Ecol. Prog. Ser. 444, 117–132.

Downs, C.A., Fauth J.E., Halas, J.C., Dustan, P., Bemiss, J. and Woodley, C.M. (2002). Oxidative stress and seasonal coral bleaching. Free Radic. Biol. Med. 33, 533–543.

Dubousquet, V., Gros, E., Berteaux-Lecellier, V., Viguier, B., Raharivelomanana, P., Bertrand, C. and Lecellier, G. (2016). Changes in fatty acid composition in the giant clam *Tridacna maxima* in response to thermal stress. Biol. Open. 5, 1400–1407.

Elfwing, T., Blidberg, E. and Tedengren, M. (2002). Physiological responses to copper in giant clams: a comparison of two methods in revealing effects on photosynthesis in zooxanthellae. Mar. Environ. Res. 54, 147–155.

Fautin, D. and Buddemeier, R. (2004). Adaptive bleaching: a general phenomenon. Hydrobiologia 530, 459–467.

Ferrier-Pages, C., Richard, C., Forcioli, D., Allemand, D., Pichon, M. and Shick, J.M. (2007). Effects of temperature and UV radiation increases on the photosynthetic efficiency in four scleractinian coral species. Biol. Bulletin. 213, 76–87.

Fitt, W.K., Brown, B.E., Warner, M.E. and Dunne, R.P. (2001). Coral bleaching: interpretation of thermal tolerance limits and thermal thresholds in tropical corals. Coral Reefs 20, 51–65.

Fitzer, S. C., Phoenix, V.R., Cusack, M. and Kamenos, N. A. (2014). Ocean acidification impacts mussel control on biomineralisation. Sci. Reports 4, 6218.

Furla, P., Galgani, I., Durand, I. and Allemand, D. (2000). Sources and mechanisms of inorganic carbon transport for coral calcification and photosynthesis. J. Exp. Biol. 203, 3445–3457.

Gazeau, F., Gattuso, J., Dawber, C., Pronker, A., Peene, F., Peene, J., Heip, C. and Middelburg, J. (2010) Effect of ocean acidification on the early life stages of the blue mussel *Mytilus edulis*. Biogeosciences 7, 2051–2060.

Gazeau, F., Parker, L.M., Comeau, S., Gattuso, J., O’Connor, W., Martin, S., Portner, H. and Ross, P.M. (2013). Impacts of ocean acidification on marine shelled molluscs. Mar. Biol. 160, 2207–2245.

Goreau, T.F. (1959). The physiology of skeleton formation in corals. I. A method for measuring the rate of calcium deposition by corals under different conditions. Biol. Bull. 116, 59–75.

Hawkins, A.J.S. and Klumpp, D.W. (1995) Nutrition of the giant clam *Tridacna gigas* (L.). II. Relative contributions of filter-feeding and the ammonium-nitrogen acquired and recycled by symbiotic alga towards total nitrogen requirements for tissue growth and metabolism. J. Exp. Mar. Biol. Ecol. 190, 263–290.

Harper, E.M. (2000). Are calcitic layers an effective adaptation against shell dissolution in the Bivalvia? J. Zool. 251, 179e186.

Hoegh-Guldberg, O., Mumby, P.J., Hooten, A.J., Steneck, R.S., Greenfield, P., Gomez, E., Harvell, C.D., Sale, P.F., Edwards, A.J., Caldeira, et al. (2007). Coral reefs under rapid climate change and ocean acidification. Science 318, 1737–1742.

Holcomb, M., Venn, A., Tambutté, E., Tambutté, S., Allemand, D., Trotter, J. and McCulloch, M. (2014). Coral calcifying fluid pH dictates response to ocean acidification. Sci. Reports 4, 5207.

Holt, A.L., Vahidinia, S., Gagnon, Y.L., Morse, D.E. and Sweeney, A.M. (2014). Photosymbiotic giant clams are transformers of solar flux. J. Royal Soc. Interface 11, 20140678.

Ikeda, S., Yamashita, H., Kondo, S., Inoue, K., Morishima, S., Koike, K. (2017). Zooxanthellal genetic varieties in giant clams are partially determined by species-intrinsic and growth-related characteristics. PLoS One 12, e0172285.

Ip, Y.K., Loong, A.M., Hiong, K.C., Wong, W.P., Chew, S.F., Reddy, K., Sivaloganathan, B. and Ballantyne, J.S. (2006). Light induces an increase in the pH of and a decrease in the ammonia concentration in the extrapallial fluid of the giant clam *Tridacna squamosa*. Physiol. Biochem. Zool. 79, 656–664.

Ip, Y.K., Hiong, K.C., Goh, E.J.K., Boo, M.V., Choo, Y.L., Ching, B., Wong, W.P. and Chew, S.F. (2017). The whitish inner mantle of the giant clam, *Tridacna squamosa*, expresses an apical Plasma Membrane Ca2+-ATPase (PMCA) which displays light-dependent gene and protein expressions. Front. Physiol. 8, 781.

IPCC (2014). Summary for policymakers. In Climate Change 2014: Impacts, Adaptation, and Vulnerability. Part A: Global and Sectoral Aspects. Contribution of Working Group II to the Fifth Assessment Report of the Intergovernmental Panel on Climate Change (ed. C.B. Field, V.R. Barros, D.J. Dokken, K.J. Mach, M.D. Mastrandrea, T.E. Bilir, M. Chatterjee, K.L. Ebi, Y.O. Estrada, R.C. Genova, B. Girma, E.S. Kissel, A.N. Levy, S. MacCracken, P.R. Mastrandrea and L.L. White), pp. 1–32. Cambridge, UK and New York, NY, USA: Cambridge University Press.

IPCC (2018) Summary for Policymakers. In Global warming of 1.5°C. An IPCC Special Report on the impacts of global warming of 1.5°C above pre-industrial levels and related global greenhouse gas emission pathways, in the context of strengthening the global response to the threat of climate change, sustainable development, and efforts to eradicate poverty. (ed. V. Masson-Delmotte, P. Zhai, H. O. Pörtner, D. Roberts, J. Skea, P. R. Shukla, A. Pirani, W. Moufouma-Okia, C. Péan, R. Pidcock, S. Connors, J. B. R. Matthews, Y. Chen, X. Zhou, M. I. Gomis, E. Lonnoy, T. Maycock, M. Tignor, T. Waterfield), pp. 1–32. World Meteorological Organization, Geneva, Switzerland.

Ishikura, M., Adachi, K. and Maruyama, T. (1999). Zooxanthellae release glucose in the tissue of a giant clam, *Tridacna crocea*. Mar. Biol. 133, 665–673.

Jones, R.J. and Hoegh-Guldberg, O. (2001). Diurnal changes in the photochemical efficiency of the symbiotic dinoflagellates (*Dinophyceae*) of corals: photoprotection, photoinactivation and the relationship to coral bleaching. Plant Cell and Environ. 24, 89–99.

Junchompoo, C., Sinrapasan, N., Penpain, C. and Patsorn, P. (2013). Changing seawater temperature effects on giant clams bleaching, Mannai Island, Rayong province, Thailand. Proceedings of the Design Symposium on Conservation of Ecosystem (The 12th SEASTAR2000 workshop), 71–76.

Kleypas, J. A. (1999). Geochemical Consequences of Increased Atmospheric Carbon Dioxide on Coral Reefs. Science 284, 118–120.

Klumpp, D.W., Bayne, B.L. and Hawkins, A.J.S. (1992). Nutrition of the giant clam *Tridacna gigas*. Contribution of filter feeding and photosynthates to respiration and growth. J. Exp. Mar. Biol. Ecol. 155, 105–122.

Klumpp, D.W. and Griffiths, C.L. (1994). Contributions of phototrophic and heterotrophic nutrition to the metabolic and growth requirements of four species of giant clam (*Tridacnidae*). Mar. Ecol. Prog. Ser. 115, 103–115.

Klumpp, D.W. and Lucas, J.S. (1994). Nutritional ecology of the giant clams *Tridacna tevoroa* and *T. derasa* from Tonga – Influence of light on filter-feeding and photosynthesis. Mar. Ecol. Prog. Ser. 107, 147–156.

Kroeker, K.J., Kordas, R.L., Crim, R.N. and Singh, G.G. (2010). Meta-analysis reveals negative yet variable effects of ocean acidification on marine organisms. Ecol. Letters. 13, 1419–1434.

Kurihara, H. (2008). Effects of CO_2_-driven ocean acidification on the early developmental stages of invertebrates. Mar. Ecol. Prog. Ser. 373, 275–284.

Kurihara, T., Yamada, H., Inoue, K., Iwai, K. and Hatta, M. (2013). Impediment to symbiosis establishment between giant clams and *Symbiodinium* algae due to sterilization of seawater. PLoS One 8, e61156.

LaJeunesse, T. C., Parkinson, J. E., Gabrielson, P. W., Jeong, H. J., Reimer, J. D., Voolstra, C. R., and Santos, S. R. (2018). Systematic revision of Symbiodiniaceae highlights the antiquity and diversity of coral endosymbionts. Current Biology 28, 2570–2580.e6.

Latchère, O., le Moullac, G., Gaetner-Mazouni, N., Fievet, J., Magré, K. and Saulnier, D. (2017). Influence of preoperative food and temperature conditions on pearl biogenesis in *Pinctada margaritifera*. Aquaculture 479, 176–187.

Leggat, W., Buck, B.H., Grice, A. and Yellowlees, D. (2003). The impact of bleaching on the metabolic contribution of dinoflagellate symbionts to their giant clam host. Plant Cell Environ. 26, 1951–1961.

Le Moullac, G., Soyez, C., Latchere, O., Vidal-Dupiol, J., Fremery, J., Saulnier, D., Lo Yat, A., Belliard, C., Mazouni-Gaertner N., and Gueguen, Y. (2016a). *Pinctada margaritifera* responses to temperature and pH: acclimation capabilities and physiological limits. Estuar. Coast Shelf Sci. 182, 261–269.

Le Moullac, G., Soyez C., Vidal-Dupiol, J., Belliard, C., Fievet, J. Sham-Koua, M., Lo Yat, A., Saulnier, D., Gaertner-Mazouni, N. et al. (2016b). Impact of pCO_2_ on the energy, reproduction and growth of the shell of the pearl oyster *Pinctada margaritifera*. Estuar. Coast Shelf Sci. 182, 274–282.

Lesser, M.P. (1996). Elevated temperatures and ultraviolet radiation cause oxidative stress and inhibit photosynthesis in symbiotic dinoflagellates. Limnol. Oceanog. 41, 271–283.

Lesser, M.P., Stochaj, W.R., Tapley, D.W. and Schick, J.M. (1990). Bleaching in coral reef anthozoans: Effects of irradiance, ultraviolet radiation and temperature on the activities of protective enzymes against active oxygen. Coral Reefs 8, 225–232.

Lim, S.S.Q., Huang, D., Soong, K., and Neo, M.L. (2019). Diversity of endosymbiotic Symbiodiniaceae in giant clams at Dongsha Atoll, northern South China Sea. Symbiosis Doi:10.1007/s13199-019-00615-5.

Lin, A.Y.M., Meyers, M.A. and Vecchio, K.S. (2006). Mechanical properties and structure of *Strombus gigas, Tridacna gigas*, and *Haliotis rufescens* sea shells: a comparative study. Mater. Sci. Eng. A. Struct. Mater. 26, 1380–1389.

Liu, W., Huang, X., Lin, J. and He, M. (2012). Seawater acidification and elevated temperature affect gene expression patterns of the pearl oyster *Pinctada fucata*. PLoS One 7, e33679.

Lucas, J.S. (1994). The biology, exploitation, and mariculture of giant clams (*Tridacnidae*). Rev. Fisheries Science 2, 181–223.

Marin, F., Le Roy, N. and Marie, B. (2012). Formation and mineralization of mollusk shell. Front. Biosci. 4, 1099–1125.

McCulloch, M., Falter, J., Trotter, J. and Montagna, P. (2012). Coral resilience to ocean acidification and global warming through pH up-regulation. Nature Clim. Change 2, 623–627.

Melzner, F., Stange, P., Trübenbach, K., Thomsen, J., Casties, I., Panknin, U., Gorb, S. and Gutowska, M. (2011) Food Supply and Seawater pCO_2_ Impact Calcification and Internal Shell Dissolution in the Blue Mussel *Mytilus edulis*. PLoS One 6, e24223.

Mies, M., Dor, P., Güth, A.Z. and Sumida, P.Y.G. (2017). Production in giant clam aquaculture: Trends and Challenges. Rev. Fish. Sci. Aquac. 25, 286–296.

Milano, S., Schöne, B.R., Wang, S. and Müller, W.E. (2016). Impact of high pCO_2_ on shell structure of the bivalve *Cerastoderma edule*. Mar. Envir. Res. 119, 144–155.

Muscatine, L., Goiran, C., Land, L., Jaubert, J., Cuif, J.P. and Allemand, D. (2005). Stable isotopes (delta C-13 and delta N-15) of organic matrix from coral skeleton. Proc. Nation. Acad. Sci. USA. 102, 1525–1530.

Neo, M.L., Eckman, W., Vicentuan, K., Teo, S.L.M. and Todd, P.A. (2015). The ecological significance of giant clams in coral reef ecosystems. Biol. Conserv. 181, 111–123.

Norton, J.H., Shepherd, M.A., Long, H.M. and Fitt, W.K. (1992). The zooxanthellal tubular system in the giant clam. Biol Bull. 183, 503–506.

Orr, J.C., Fabry, V.J., Aumont, O., Bopp, L., Doney, S.C., Feely, R.A., Gnanadesikan, A., Gruber, N., Ishida, A., Joos, F. et al. (2005). Anthropogenic ocean acidification over the twenty-first century and its impact on calcifying organisms. Nature 437, 681–686.

Palmer, A.R. (1992). Calcification in marine molluscs: how costly is it? Proc. Natl. Acad. Sci. USA. 89, 1379–1382.

Pätzold, J., Heinrichs, J.P., Wolschendorf, K. and Wefer, G. (1991). Correlation of stable oxygen isotope temperature record with light attenuation profiles in reef-dwelling *Tridacna* shells. Coral Reefs 10, 65–69.

Pearse, V.B. and Muscatine, L. (1971). Role of symbiotic algae (zooxanthellae) in coral calcification. Biol. Bull. 141, 350–363.

Perez, S. and Weis, V. (2006). Nitric oxide and cnidarian bleaching: an eviction notice mediates breakdown of a symbiosis. J. Exp. Biol. 209, 2804–2810.

Pinzón, J.H., Devlin-Durante, M.K., Weber, M.X., Baums, I.B., and LaJeunesse, T.C. (2011). Microsatellite loci for *Symbiodinium* A3 (*S. fitti*) a common algal symbiont among Caribbean *Acropora* (stony corals) and Indo-Pacific giant clams (*Tridacna*). Conserv. Genet. Resour. 3, 45–47.

Richier, S., Sabourault, C., Courtiade, J., Zucchini, N., Allemand, D. and Furla, P. (2006). Oxidative stress and apoptotic events during thermal stress in the symbiotic sea anemone, *Anemonia viridis*. Febs Journal 273, 4186–4198.

Richter C, Rühle, W. and Wild, A. (1990). Studies on the mechanisms of photosystem II photoinhibition II. The involvement of toxic oxygen species. Photosynthesis Res. 24, 237–243.

Ries, J.B., Cohen, A.L. and McCorkle, D.C. (2009). Marine calcifiers exhibit mixed responses to CO_2_-induced ocean acidification. Geology 37, 1131–1134.

Ries, J.B. (2011) A physiochemical framework for interpreting the biological calcification response to CO_2_-induced ocean acidification. Geochim Cosmochim Acta 75, 4053–4064.

Rodolfo-Metalpa, R., Houlbreque, F., Tambutte, E., Boisson, F., Baggini, C., Patti, F.P., Jeffree, R., Fine, M., Foggo, A., Gattuso, J.P. et al. (2011). Coral and mollusc resistance to ocean acidification adversely affected by warming. Nature Clim. Change 1, 308–312.

Sabine, C.L., Feely, R.A., Gruber, N., Key, R.M., Lee, K., Bullister, J.L., Wanninkhof, R., Wong, C.S., Wallace, D.W.R., Tilbrook, B. et al. (2004). The oceanic sink for anthropogenic CO_2_. Science 305, 367–371.

Schwartzmann, C., Durrieu, M., Sow, M., Ciret, P., Lazareth, C.E. and Massabuau, J.-C. (2011) In situ giant clam growth rate behavior in relation to temperature: A one-year couple study of high-frequency noninvasive valvometry and scleraochronology. Limnol. Ocean. 56, 1940–1951.

Smith, D.J., Suggett, D.J. and Baker, N.R. (2005). Is photoinhibition of zooxanthellae photosynthesis the primary cause of thermal bleaching in corals? Global Change Biology 11, 1–11.

Soo, P. and Todd, P.A. (2014). The behaviour of giant clams. Mar. Biol. 161, 2699–2717.

Talmage, S.C. and Gobler, C.J. (2011). Effects of elevated temperature and carbon dioxide on the growth and survival of larvae and juveniles of three species of northwest Atlantic bivalves. PLoS One 6, e26941.

Toonen, R.J., Nakayama, T., Ogawa, T., Rossiter, A. and Delbeek, J.C. (2012). Growth of cultured giant clams (*Tridacna spp.*) in low pH, high nutrient seawater: species-specific effects of substrate and supplemental feeding under acidification. J. Mar. Biol. Assoc. UK. 92, 731–740.

Vandermeulen, J.H., Davis, N.D. and Muscatine, L. (1972). The effect of inhibitors of photosynthesis on zooxanthellae in corals and other marine invertebrates. Mar. Bio. 16, 185–191.

van Heuven, S., Pierrot, D., Lewis, E. and Wallace, D.W.R. (2009). MATLAB Program developed for CO_2_ system calculations, ORNL/CDIAC-105b, Carbon Dioxide Information Analysis Center, Oak Ridge National Laboratory, US Department of Energy, Oak Ridge, Tennessee.

Van Wynsberge, S., Andréfouët, S., Gaertner-Mazouni, N., Wabnitz, C.C.C., Gilbert, A., Remoissenet, G., Payri, C. and Fauvelot, C. (2016). Drivers of density for the exploited giant clam *Tridacna maxima*: a meta-analysis. Fish Fish. 17, 567–584.

Venn, A., Tambutté, E., Holcomb, M., Laurent, J. Allemand, D. and Tambutté, S. (2013). Impact of seawater acidification on pH at the tissue-skeleton interface and calcification in reef corals. PNAS 110, 1634–1639.

Warner, M.E., Fitt, W.K., and Schmidt, G.W. (1999). Damage to photosystem II in symbiotic dinoflagellates: A determinant of coral bleaching. Proc. Natl. Acad. Sci. 96, 8007–8012.

Waters, C.G. (2008). Biological responses of juvenile *Tridacna maxima* (Mollusca:Bivalvia) to increased pCO_2_ and ocean acidification. Master thesis, The Evergreen State College.

Watson, S.-A., Southgate, P.C., Miller, G.M., Moorhead, J.A. and Knauer, J. (2012). Ocean acidification and warming reduce juvenile survival of the fluted giant clam *Tridacna squamosa*. Molluscan Res. 32, 177–180.

Watson, S. (2015). Giant Clams and rising CO_2_: Light may ameliorate effects of ocean acidification on a solar-powered animal. PLoS One 10, e0128405.

Welladsen, H.M., Southgate, P.C. and Heimann, K. (2010). The effects of exposure to near-future levels of ocean acidification on shell characteristics of *Pinctada fucata* (Bivalvia: Pteriidae). Molluscan Res. 30, 125–130.

Yau, A.J.Y. and Fan, T.Y. (2012). Size-dependent photosynthetic performance in the giant clam *Tridacna maxima*, a mixotrophic marine bivalve. Mar. Biol. 159, 65–75.

Zeebe, R.E., Zachos, J.C., Caldeira, K. and Tyrrell, T. (2008). Oceans-Carbon emissions and acidification. Science 321, 51–52.

Zhou, Z., Liu, Z., Wang, L., Luo, J. and Li, H. (2019) Oxidative stress, apoptosis activation and symbiosis disruption in giant clam *Tridacna crocea* under high temperature. Fish. Fish. Immunol. 84, 451–457.

